# Convergent pathways of reductive mitochondrial evolution characterised with hypercubic inference

**DOI:** 10.1101/2025.04.29.651065

**Authors:** Robert C. Glastad, Iain G. Johnston

## Abstract

For a striking example of mitochondrial behaviour beyond ATP generation, consider mitochondrion-related organelles (MROs). Hydrogenosomes, mitosomes, and other reduced mitochondrial forms have evolved through the loss of physical and functional features, from individual ETC complexes to oxidative phosphorylaytion and the very ability to produce ATP (and further). Reduction of mitochondria is a dramatic example of convergent evolution, occuring in every eukaryotic kingdom and many parallel times. Here, we use hypercubic inference, a class of methods from evolutionary accumulation modelling (EvAM), to explore the pathways of convergent mitochondrial reduction across eukaryotes. We find that most MRO diversity can be explained by small variations on two distinct pathways, starting with either the loss of Complex I or the loss of Complexes III/IV, which tend to proceed over different characteristic timescales. We show that different clades, including ciliates and apicomplexans, reflect particular instances of these pathways. Using metabolic modelling, we connect the structure of these evolutionary pathways to the metabolic impact of the changes involved, suggesting a plausible explanation for the dramatically convergent nature of reductive evolution. We discuss this approach in connection with related theory on the genetic and functional reduction of mitochondria across organisms.

## Introduction

Mitochondria are central, multifunctional organelles in eukaryotic cells (Monzel et al., 2023). Originally independent organisms, over evolutionary history they have become progressively entrained by their “host” cells, with most of their genes having been lost or transferred to the host nucleus (Giannakis et al., 2022; Janouškovec et al., 2017; Johnston & Williams, 2016; Roger et al., 2017; Smith & Keeling, 2015; Veeraragavan et al., 2024). In some species – including well-known cases in yeasts (Hackstein et al., 2006), parasitic plants (Petersen et al., 2020; Senkler et al., 2018), and anaerobic protists (Leger et al., 2019; Makiuchi & Nozaki, 2014; Mathur et al., 2021) – this genetic reduction has been accompanied by the loss of physical and functional features of mitochondria (Maciszewski & Karnkowska, 2019; Makiuchi & Nozaki, 2014; Stairs et al., 2015). These features include electron transport chain (ETC) protein complexes, transporters, the mitochondrial genome, elements of the tricarboxylic acid (TCA) cycle, and even the ability to produce ATP. The collection of species with observed mitochondrial feature losses continues to expand, including discoveries of the first eukaryote lacking mitochondria (Karnkowska et al., 2016), the first animal lacking mtDNA (Yahalomi et al., 2020), and increasing characterisation of reduced mitochondria in protists (Mathur et al., 2021; Záhonová et al., 2023).

These highly reduced organelles can generally be referred to as mitochondrion-related organelles or MROs (de Paula et al., 2012; Hampl & Roger, 2024; Leger et al., 2019; Lewis et al., 2020; Makiuchi & Nozaki, 2014; Van Der Giezen, 2009). They include hydrogenosomes (which retain the ability to produce ATP, including through a characteristic pathway producing molecular hydrogen), mitosomes (which are highly reduced and do not produce ATP) and other categories (sometimes labelled class I-V dependent on metabolic status (Müller et al., 2012)). Reduced mitochondria and MROs retain metabolic importance in their carrier species (notably, machinery involved in iron-sulfur cluster assembly seems to be near-universally retained in MROs), with some remaining features postulated as targets for therapeutic interventions (Goodman et al., 2017). But the loss of canonical mitochondrial features often leads to unusual behaviour in these MROs. In the absence of a complete proton-pumping electron transport chain (ETC), ATP synthase may run in reverse, consuming ATP to maintain mitochondrial membrane potential. Genes for mitochondrial machinery may be retained without lab-detectable activity of that machinery (Gerasimov et al., 2025; Verner et al., 2011). Loss of proton-pumping ETC complexes may be partly compensated by non-proton-pumping alternatives, including alternative NADH dehydrogenases and alternative oxidases (Fang & Beattie, 2003; Mathur et al., 2021; Rasmusson et al., 2008). Clearly, these reductions highlight roles of mitochondria(−related organelles) beyond ATP production.

MROs have been found in diverse and deeply-branching eukaryotic lineages, making the loss of mitochondrial features a striking example of convergent evolution (Lewis et al., 2020; Mathur et al., 2021; Stairs et al., 2015; Van Der Giezen, 2009). As such, similarities and differences in the “pathways” of the loss process across different clades – the orderings and timings with which features are lost – are points of fundamental biological interest. Algorithmic developments in the study of convergent evolutionary processes have made it possible to quantify structure, variability, and uncertainty in the pathways by which such processes take place (Aga et al., 2024; Beerenwinkel et al., 2005; Boyko & Beaulieu, 2021, 2023; Diaz-Uriarte & Herrera-Nieto, 2022; Renz et al., 2024; Williams et al., 2013).

While these approaches have been applied to organelle genome evolution (Giannakis et al., 2022; Giannakis, Aga, et al., 2024; Johnston & Williams, 2016), they have not to our knowledge yet been applied to this class of reductive mitochondrial evolution. Here, we aim to apply these “hypercubic inference” approaches – a class of methods within evolutionary accumulation modelling or EvAM (Diaz-Uriarte & Herrera-Nieto, 2022; Diaz-Uriarte & Johnston, 2024) -- to describe the possible evolutionary pathways of mitochondrial feature loss across eukaryotes.

## Methods

### Dataset compilation

We required a set of features that could (when required) be estimated from genetic information, that have previously been well studied, that reflect different aspects of mitochondrial functionality, and which could be captured by simple (and therefore generalisable) metabolic modelling approaches. Motivated by an extensive literature on the key features categorising mitochondrial reduction, the features we consider (with shorthand labels) are: the TCA cycle (*TCA*), ETC respiratory protein complexes I-IV (*CI*; *CII*; *CIII*; *CIV*), ATP synthase (*CV*), the mtDNA genome (*DNA*), pyruvate dehydrogenase (*PDH*), and machinery required for iron-sulfur cluster production (*Fe-S*). We took advantage of the extensive literature on the topic to compile lists from meta-analysis of MROs as well as a collection of individual species where these features were characterised (reference list in Supp. Text 1). The data, with sources and discussion, is available at the Github repository below.

### Binarisation

To assign binary presence-absence markers to these sometimes nuanced features, we make several feature-specific choices. For ETC complexes, we consider “presence” to mean the canonical versions of complexes found in most eukaryotes. Some yeasts and protists have replaced canonical CI with an alternative NADH dehydrogenase; this counts as loss in our protocol. Some organisms have replaced CIV (and CIII) with alternative oxidases as terminal electron acceptors and/or replaced PDH with a non-canonical alternative (Mathur et al., 2021); we also count these as losses. For the TCA cycle, we consider the complete, common TCA cycle as “presence” and incomplete cycles as “absence”.

### Tree construction

We used both NCBI’s Common Taxonomy Tree (Federhen, 2012) to obtain a coarse-grained taxonomy linking species, and TimeTree (Kumar et al., 2022) for the subset of available species for a smaller phylogeny with estimated timings.

### Hypercubic transition path sampling (HyperTraPS)

HyperTraPS considers how multiple, potentially coupled, binary features coevolve on a phylogeny (Aga et al., 2024; Greenbury et al., 2020; Johnston & Williams, 2016). Given a phylogeny where each tip has a collection of *L* binary features (for example, Fig. 1A), the approach first reconstructs ancestral states through the phylogeny. Here, we assume that mitochondrial feature losses are relatively rare and irreversible, so that an ancestor is inferred to have lost a feature if and only if all its descendants have lost it (otherwise, loss is assumed to occur independently in the subset of descendant lineages lacking the feature). Each ancestor-descendant pair on the phylogeny then constitutes an observed “before-after” pair of states (which may be identical), and a timescale (if the phylogeny has meaningful branch lengths).

**Figure 1.**
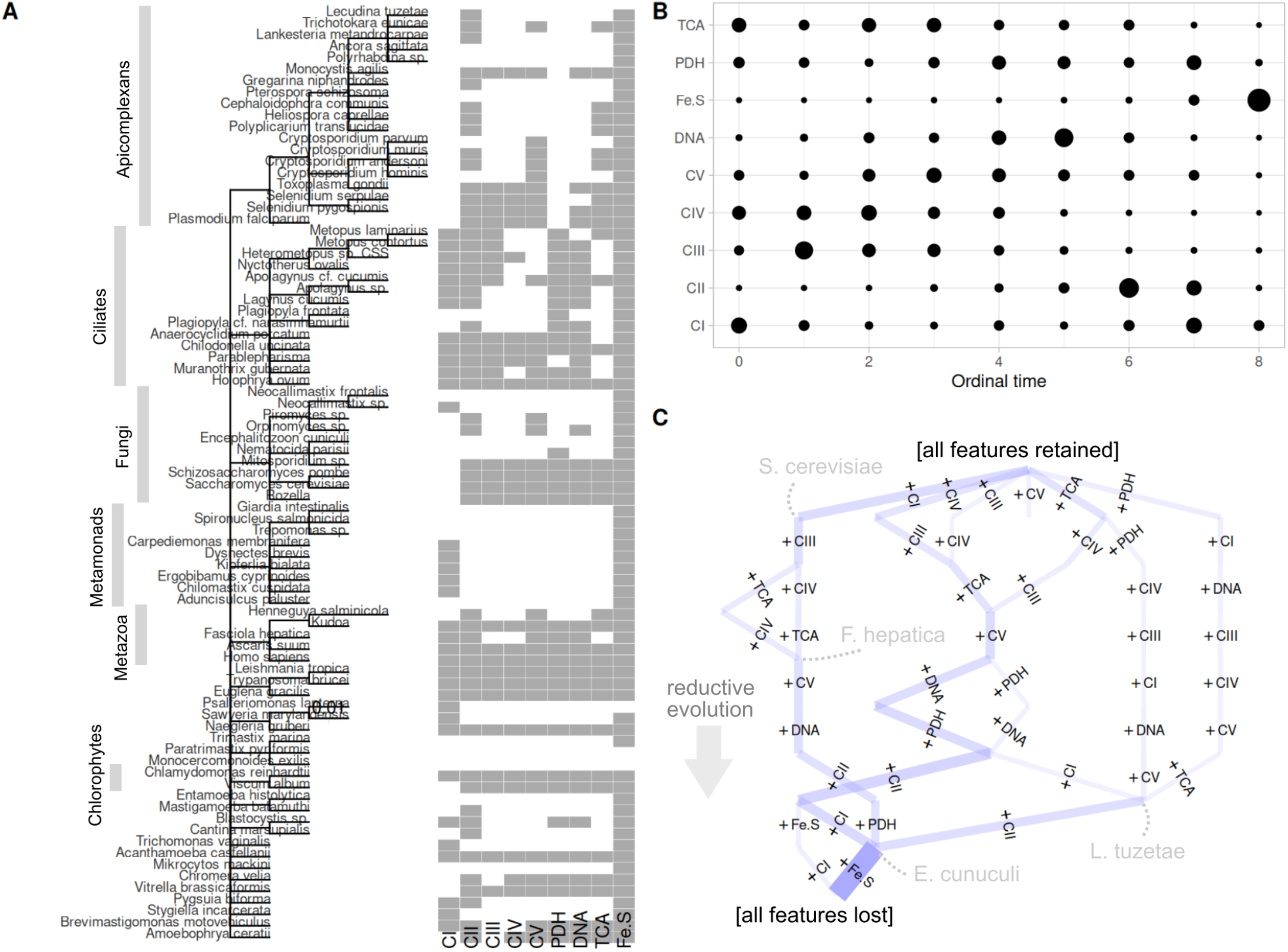
Data and inference for pathways of reductive mitochondrial evolution across eukaryotes. **(A)** Source data, comprising the taxonomy of species and the presence (grey) or absence (white) of each mitochondrial feature in each species. Selected clades are labelled. **(B)** “Bubble plot” from HyperTraPS inference. The area of each circle gives the probability with which the corresponding feature (rows) is lost at the corresponding ordering (columns) in a reductive evolution pathway. For example, *Fe-S* is overwhelmingly likely to be the last feature lost; *CI* has two peaks in loss probability, as the first or eigth feature to be lost. **(C)** The most likely transitions through the evolutionary space inferred by HyperTraPS. Each edge (line segment) corresponds to the loss of a feature, with width giving the probability of that loss; each node (point between edges) corresponds to a particular set of presences and absences. Nodes corresponding to some example species are labelled. The topmost state is the ancestral state with all features present; the bottommost state has no features remaining; reductive evolution proceeds down the figure. For example, the state at the far left of the figure has lost *CI, CIII*, and *TCA*, and its most likely next loss is *CIV*.

HyperTraPS works by considering a “state space” of all the 2^*L*^ possible binary states that could, in principle, be observed in an evolutionary system. Transitions through this space are allowed from a “before” state to an “after” state that differs from the before state by the loss of exactly one feature. All possible paths from the state 111… (retaining all features) to the state 000… (having lost all features) are therefore possible. Each transition is parameterised by a probability, or a rate (in the continuous time case). HyperTraPS then identifies parameters for this space of evolutionary transitions that are most compatible with the set of “before-after” pairs of states observed in the dataset. By default (and here), this is a Bayesian inference process using Markov chain Monte Carlo (Aga et al., 2024). After checking for numerical convergence, the results can be summarised in various ways, including the probability that a given feature is the *n*th to be lost in the reduction process (Fig. 1B) or a summary of the specific inferred transition parameters in the evolutionary space (Fig. 1C).

By default (and here), HyperTraPS considers parameters describing (a) the base rate of loss of each feature and (b) how this base rate is affected by the loss of each other feature (Johnston & Williams, 2016) – a scheme also known as mutual hazard networks (Schill et al., 2020). In this way, interactions between feature losses – whether the loss of *A* makes the loss of *B* more or less likely – can also be quantified.

### Flux balance analysis

We used MitoMAMMAL, a recent genome-scale metabolic model of mammalian mitochondria, as our model for non-reduced mitochondrial metabolism (Chapman et al., 2025). Using *sybil* (Gelius-Dietrich et al., 2013) and *sybilSBML* (Fritzemeier et al., 2017) packages in R, we performed flux balance analysis (Orth et al., 2010) to explore influences of reaction losses on mitochondrial(−related organelle) metabolism. We set either ATP production or generation of proton-motive force as the objective function of interest, and modelled the influence of removing reactions corresponding to ETC Complexes I-V, individual TCA cycle steps, and pyrvuate dehydrogenase. We modelled metabolism both under default oxygen conditions and under hypoxic conditions, modelled with oxygen levels decreased to 10% of the default case. To model the presence of compensatory, non-canonical ETC subunits, we introduced reactions:

AOX: 2 ubiquinol + O_2_ → 2 ubiquinone + H_2_O

NDHalt: H^+^ + NADH + 0.999 ubiquinone + 0.002 O_2_ → NAD + 0.999 ubiquinol + 0.002 O_2_·^-^

These two reactions serve as an electron acceptor and an NADH reducer respectively without pumping protons. The small deviations from integers in the stoichiometry of the NDHalt reaction are taken from the original MitoMAMMAL CI reaction, and reflect some small amount of superoxide production. Superoxide production has also been suggested in alternative NADH dehydrogenases in yeast (Fang & Beattie, 2003), but the presence or absence of superoxide is not an consequential term in our metabolic analysis and will not influence our findings.

### Software

In addition to the packages referenced above, we used R (R Core Team, 2022) with *ggplot2* (Wickham, 2016), *ggpubr* (Kassambara, 2020), *ggraph* (Pedersen, 2020), *ggtree* and *ggtreeExtra* (Xu et al., 2021; Yu et al., 2017) for visualisation; *igraph* (Csardi & Nepusz, 2006) for network analysis; and *ape* (Paradis & Schliep, 2019), *phangorn* (Schliep, 2011), and *phytools* (Revell, 2012) for phylogenetics. The specific parameters used for the HyperTraPS algorithm were 200 sampling walkers for likelihood calculations, 10^5^ MCMC steps, no likelihood penalisation, and a normal perturbation kernel with standard deviation 0.5, checking for convergence across three numerical replicates (Aga et al., 2024). Our pipeline and dataset is freely available at https://github.com/StochasticBiology/MROs.

## Results

### Distinct pathways of mitochondrial feature loss in reductive evolution across eukaryotes

We first sought to compile a dataset of patterns of lost mitochondrial features across eukaryotes (Supp. Data 1). This process involved (a) selecting the mitochondrial and MRO features that we would characterise; (b) determining a protocol for labelling each feature “present” or “absent” in each species; (c) compiling a list of species and their presence-absence patterns; and (d) determining the relationship between these species. Details of these steps are given in the Methods. To summarise, we consider losses of canonical ETC complexes (*CI*-*CV*), loss of any component of the canonical TCA cycle (*TCA*), loss of mtDNA (*DNA*), loss of canonical pyruvate dehydrogenase (*PDH*), and loss of iron-sulfur metabolism capacity (*Fe-S*), across a set of eukaryotes connected by an estimated taxonomy (Fig. 1A) and a subset with a better time-resolved phylogeny (Supp. Fig. 1A). If a lost feature is replaced by a non-canonical alternative, we still consider it lost.

The datasets are shown in Fig. 1A and Supp. Fig. 1A. It should immediately be pointed out that the features we consider are not independent. As CII is a component of the complete TCA cycle, we cannot have loss of *CII* without loss of *TCA*. As essential subunits of CIII and CIV are encoded in mtDNA, we cannot have loss of *DNA* without loss of *CIII* and *CIV* – although our dataset contains one exception, *Selenidium serpulae*, where a mitochondrial genome appears to be absent and CIII-IV genes are speculated to have been transferred to the nucleus (Mathur et al., 2021).

Our first step in analysing these data was to use HyperTraPS (hypercubic transition path sampling (Aga et al., 2024; Greenbury et al., 2020; Johnston & Williams, 2016)) to infer the likely orderings of feature loss events across these trees of species (see Methods). Briefly, HyperTraPS constructs an evolutionary model where all combinations of feature presence and absence are possible, then asks which weightings of transitions through this space (from a putative ancestor with all features, towards a putative eventual “end state” with no features) are most compatible with the observed data (controlling for relatedness, in the sense that descendants of a common ancestor do not constitute independent representatives of a given set of features (Maddison & FitzJohn, 2015; Revell, 2010)). First results from this inference are shown in Fig. 1B-C and Supp. Fig. 1B-C. In Fig. 1B and Supp. Fig. 1B, the probability that a given feature is lost at a given step in reductive evolution (for example, the first feature to be lost, the second, and so on) is given. As a “control”, iron-sulfur metabolism (*Fe-S*) is always inferred to be most likely lost last. Complex I (*CI*) has a clearly bimodal loss distributions: it may be lost early, or late, in the reductive process, but less likely at intermediate stages. This immediately suggests multiple distinct evolutionary pathways: one type with early loss of *CI* and one distinct type with late loss of *CI*.

The presence of multiple distinct pathways is supported by a more detailed look at the inferred transitions through the evolutionary model (Fig. 1C, Supp. Fig. 1C). Here, the most likely transitions from the (top) ancestral state towards the (bottom) putative final state are shown. It can immediately be seen that the most likely evolutionary pathways are highly canalised: following a collection of possible first loss events, subsequent evolution follows distinct pathways before converging on *Fe-S* as the final loss event.

Of course, an important issue in any such analysis pipeline is the robustness of results to errors or nuances in the original dataset. We first note that the discrete-time and continuous-time inference processes for the different data subsets in Fig. 1 and Supp. Fig. 1 give compatible results. We also performed a control study where we artificially perturbed the source data by flipping a random 10% of the presence-absence markers for ten resampled datasets (see Methods). The resulting inferred dynamics were largely consistent with each other and with the unperturbed results (Supp. Fig. 2), suggesting that these findings are robust to (at least) this error rate in the source data.

**Figure 2.**
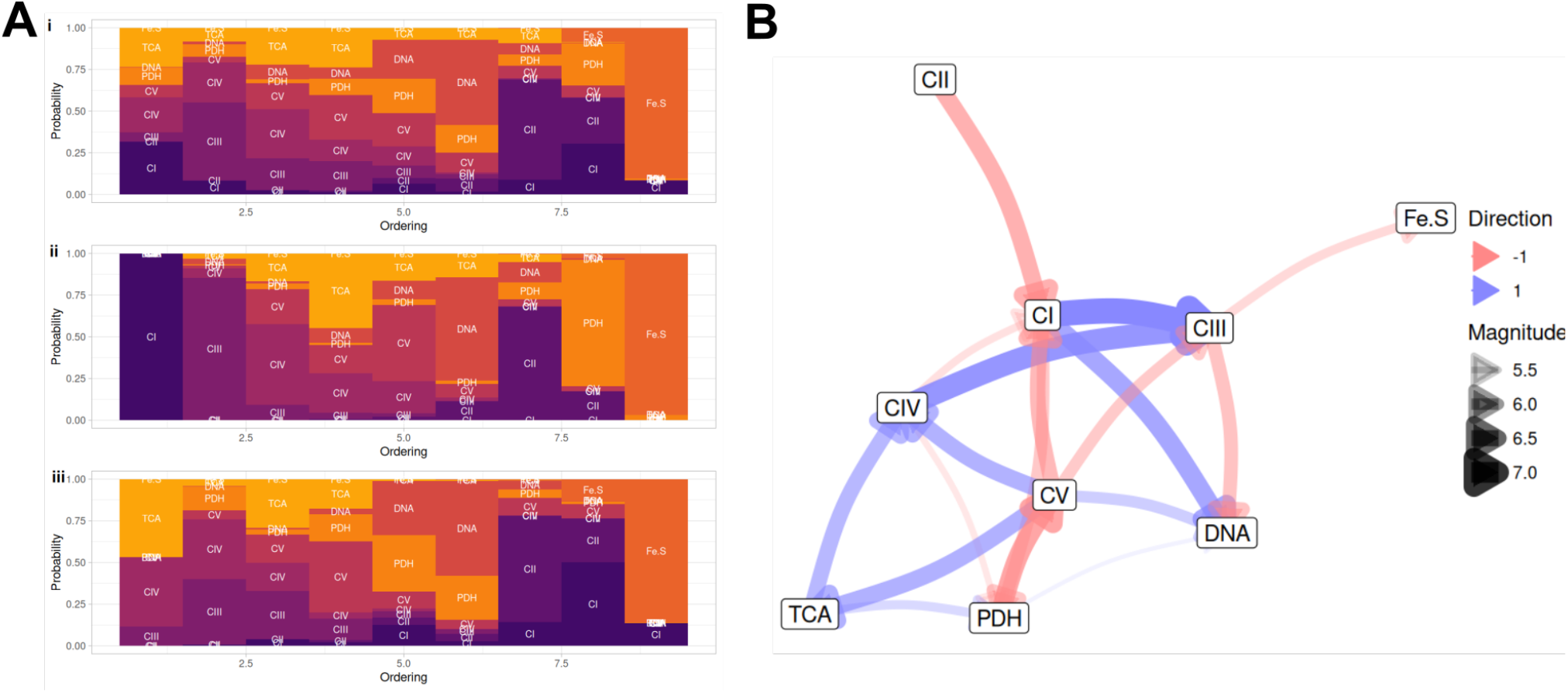
Influences between features, and evolutionary pathway structures. **(A)** “Motif plots” from HyperTraPS output describing the probability (section height) that each feature is lost at each ordering step in a pathway (the same concept as Fig. 1B). (i) All pathways, reproducing Fig. 1B. (ii) CI-class pathway, where the first loss is constrained to be *CI* and the conditional probabilities of following losses are plotted through the pathway. (iii) alt-class pathway, where the first loss is constrained to be either *CIII, CIV*, or *TCA*. **(B)** Inferred influences between feature losses; edge widths given posterior means (in effectively arbitrary units) for parameters with coefficients of variation under 0.65. Blue edges are “activating” (*CIV* → *CIII* means that *CIV* loss makes *CIII* loss more likely); red edges are “repressing” (*CIV* loss makes *CI* loss less likely).

### Inferred pathway structures and influences in reductive mitochondrial evolution

HyperTraPS makes it possible to dissect the structure of evolutionary pathways, and the possible interactions between events that support them, in more depth. In Fig. 2A, we plot the inferred probabilities with which different subsequent loss events follow a particular, given first loss event. Strikingly distinct pathways result when this given first loss event is either *CI* or an alternative. These initial steps lead to rather tightly-defined following pathways inferred to occur with high probability, which we will refer to as:

CI-class: following *CI, CIII*, (*CIV* and *TCA*), *CV, DNA, CII, PDH*, (then *Fe-S*).

alt-class: following (*CIII, CIV*, and *TCA*), *CV, DNA, PDH, CII, CI*, (then *Fe-S*).

It will immediately be seen that the combination of these distinct CI- and alt-class pathways can give rise to the bimodality in *CI* ordering distribution observed in Fig. 1B. There is also a lower-probability set of pathways involving CI- and alt-class orderings with *PDH* loss evolving earlier, rather than later.

HyperTraPS allows features to influence each other, in the sense that the loss of one feature can have a positive (activating, →) or negative (repressing, ⊣) effect on the probability that another feature is lost. Every feature has a base rate of loss, and the loss of other features can influence this base rate. Fig. 2B shows the collection of these influences that were robustly inferred across eukaryotic species. It is readily seen that positive influences *CIV* → *CIII* and *TCA* → *CIV* support the observed close links between the first steps of the alt-class pathway, and *CI* → *CIII* supports these steps immediately after the CI-class pathway begins. The *CIV* ⊣ *CI* interaction supports the distinction between the CI-class and alt-class pathways, ensuring that once the alt-class pathway starts, *CI* loss is forced later. Mutual repressions between *CV* and *PDH* support the bimodal distribution of *PDH* in the CI-class pathway.

A recent approach using hypercubic directed acyclic graphs (HyperDAGs) (Giannakis, Aga, et al., 2024) allows a minimal set of evolutionary pathways to be computed for a dataset. In a sense, the most deterministic, reproducible evolutionary system corresponds to a single, canalised evolutionary pathway in Fig. 1C. Every independent lineage would then evolve through the same sequence of loss events. Every instance where one species has *A* but not *B* and another species has *B* but not *A* (referred to as an “incompatibility”) requires departure from such a single, canalised evolutionary pathway. Such departures take the form of branches – out-degrees greater than 1 – in the evolutionary spaces like Fig. 1C. HyperDAGs finds the smallest number of branches required to account for the incompatibilities in a dataset; here only 6 branches are required to account for all 82 observations (Supp. Fig. 3). Of these, 5 are associated with different early orderings of *CIII, CIV*, and *TCA* loss (and are therefore in a sense contained within the alt-class pathway), and the final branch corresponds to whether *PDH* is acquired early or late when *CI* is the first loss.

**Figure 3.**
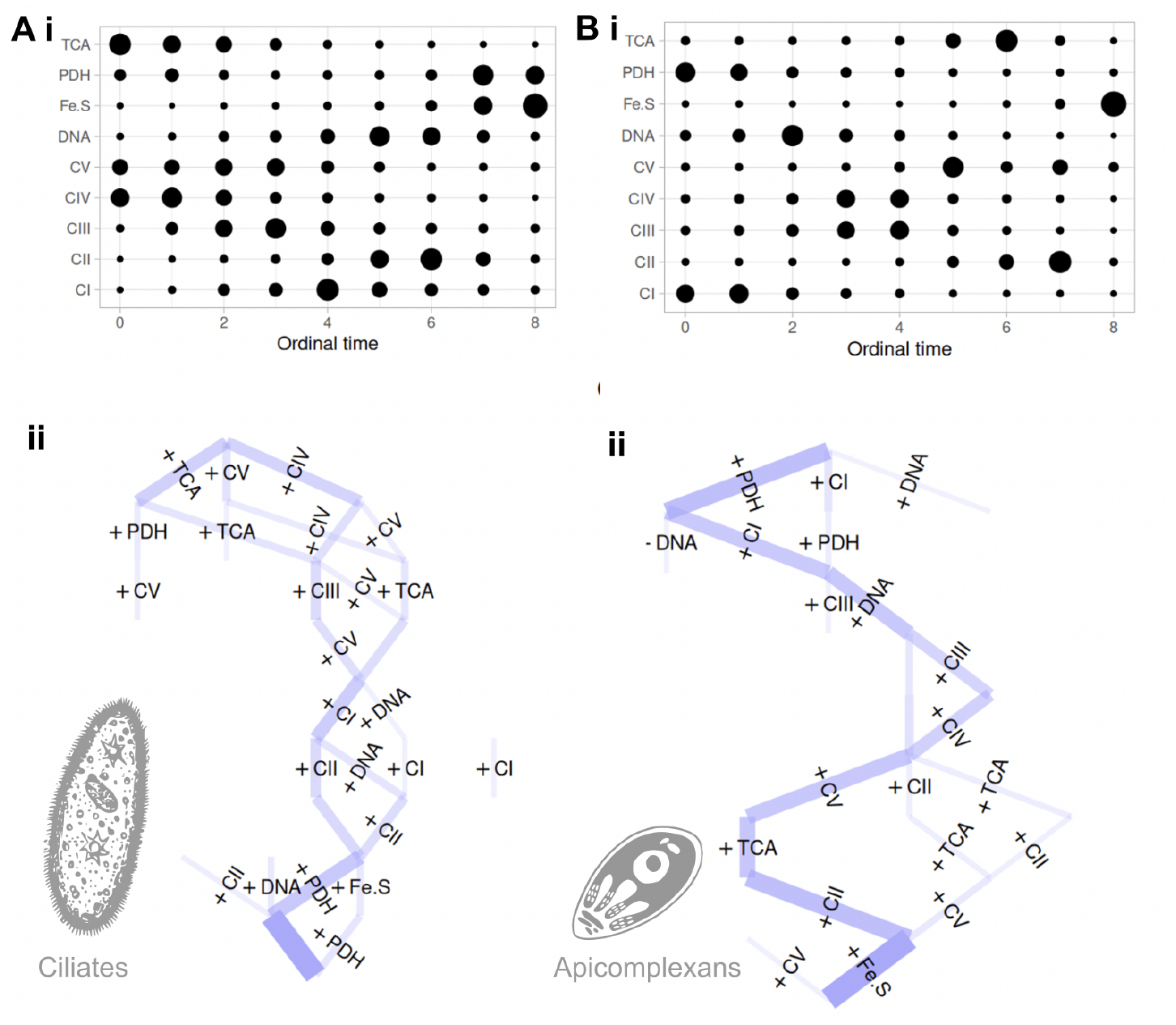
Evolutionary pathways inferred within specific protist phyla. **(A)** (i) Bubble plot (as in Fig. 1B) and (ii) transition network (as in Fig. 1C) for HyperTraPS inference using only the ciliate phylum as input data. **(B)** (i) Bubble plot and (ii) transition network using only the apicomplexan phylum as input data. Inset images are public domain illustrations from phylopic.org.

This suggestion of distinct pathways is supported by individual species profiles in Fig. 1A, including (for example): the well-known mistletoe (*Viscum album*) and yeast (*Saccharomyces cerevisiae*) reflecting early CI-class, *Nematosida parisii* and *Plagiopyla frontata* reflecting late CI-class, *Brevimastigomonas motovehiculus* and liver fluke (*Fasciola hepatica*) reflecting early alt-class, and *Neocallimastix* sp. and *Chilomastix cuspidata* reflecting late alt-class. Taken together, these results consistently suggest two distinct, canalised evolutionary pathways of reductive mitochondrial evolution.

### Clade-specific pathways of inferred reductive mitochondrial evolution

Following these observations, we next asked whether these structures in evolutionary pathways across eukaryotes arose from different behaviour in different clades. The first clade we examined was ciliates, a phylum of widespread protists (Fig. 1A, (Rotterová et al., 2020)). Here, while reduction of CI is observed via the loss of subunits, complete loss of the complex seems less common. Accordingly, HyperTraPS shows no support for the CI-class pathway in ciliates, with the alt-class pathway dominating the inferred dynamics (Fig. 3A-B).

We also explored dynamics in apicomplexans, a phylum of largely parasitic protists (Fig. 1A, (Mathur et al., 2021)). Here, CI is commonly lost, and PDH is also lost across the phylum (Fig. 3C-D). Accordingly, HyperTraPS recovers strong support for the CI-class pathway, with the change that *PDH* is lost as the first or second step rather than later. The inferred evolutionary trajectory in this phylum is very highly canalised (Fig. 3D). HyperDAG analysis shows that only one incompatibility exists (*CII* vs *CV*) so that, in principle, a single evolutionary pathway with one branch can explain all apicomplexan observations.

### Inference about timescales of reductive evolution pathways

For the subset of species with available estimates of divergence times (Supp. Fig. 1A), a version of the algorithm called HyperTraPS-CT (continuous time) allows inference about the possible timescales of evolutionary events (as opposed to only considering their relative orderings, as in Figs. 1-2. We exploit HyperTraPS-CT’s capacity to account for uncertainty in timings and assume a time window for each branch between half and twice the point estimate from TimeTree (Kumar et al., 2022). As above, the summaries of results from this continuous time inference (Supp. Fig. 1B-C) are compatible with the ordering-only case, with some shifts. First, ordering probabilities are more uniform, reflecting the increased uncertainty from the necessarily smaller dataset. The alt-class and CI-class pathways are roughly equal in probability, and *PDH* is more likely acquired early in the CI-class pathway.

The inferred timescales of loss events are illustrated in Supp. Fig. 4. The distributions of likely timings of loss events are very broad – both owing to the diversity of lineages where the losses occur and the relatively small dataset size. Despite this breadth, a general trend is visible, where the distributions of alt-class loss steps are shifted towards shorter timescales than for CI-class (though with considerable overlap). This reflects the set of species where *CI* is lost with no other losses (resembling a “stalling” of the loss process) versus the set of species where multiple losses in the *CIII, CIV, TCA* group, and followup steps in the alt-class pathway, have occurred in comparatively rapid succession. Notably, the first three steps of the alt-class pathway and the second to fourth steps of the CI-class pathway – in both cases most likely to involve *CIII, CIV, TCA* losses – occur within what is comparatively a very short timescale of each other.

**Figure 4.**
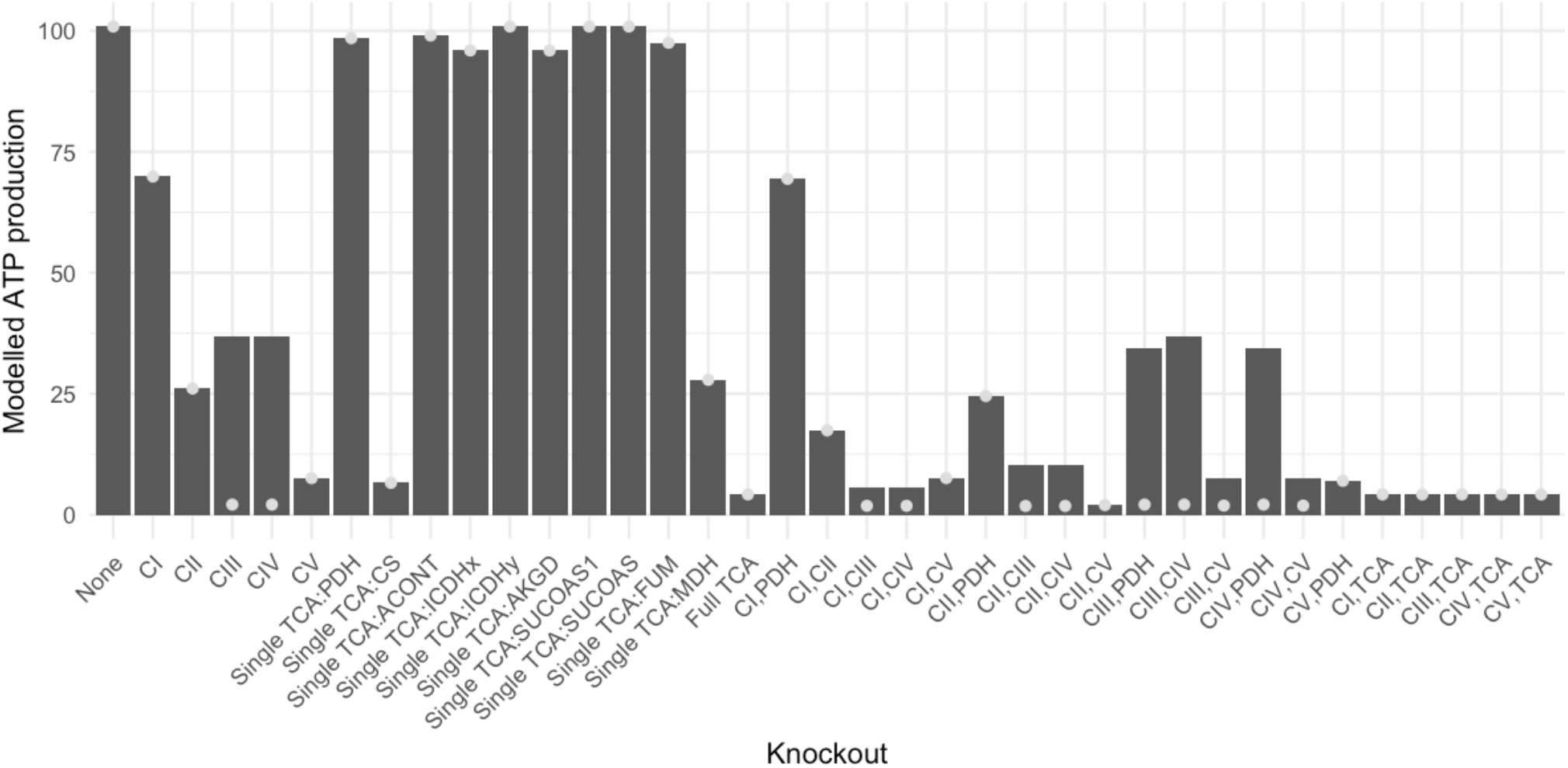
Flux balance analysis of the metabolic impact of feature loss on ATP production. The computationally predicted metabolic impact of the loss of single features and sets of features in our study, using MitoMAMMAL (Chapman et al., 2025). Each column reflects either the non-reduced case (“None”), a single computational knockout of a metabolic reaction, or a set of simultaneous knockouts. The height of a bar gives predicted ATP production under that knockout experiment. Grey circles give the predicted ATP production when alternative oxidase is not modelled as a terminal electron acceptor (see Methods). Individual TCA cycle reactions are labelled with shorthand acronyms; impactful reactions are citrate synthase (CS) and malate dehydrogenase (MDH). Other conditions and objectives are shown in Supp. Fig. 5.

### Connecting inferred loss dynamics and simulated metabolic impact of lost mitochondrial features

We next asked what the metabolic implications of these different patterns of mitochondrial feature loss could be. Working as generally as possible, we used MitoMAMMAL, a recent genome-scale model of mammalian mitochondrial metabolism (Chapman et al., 2025) (reflecting an ancestral, non-reduced organelle). We performed a constraint-based computational analysis of the metabolic impact of each loss on objective functions of ATP production and proton-motive force generation in normoxic and hypoxic conditions, including in the presence of introduced reactions reflecting non-canonical replacements of ETC complexes (see Methods).

Under normoxic conditions (Fig. 4), from the ETC complexes, the loss of *CI* has the lowest effect on ATP production, followed by the loss of *CIII* and/or *CIV* with compensatory gain of AOX. Without compensatory gain of AOX, *CIII/IV* loss immediately compromises ETC activity due to the lack of a terminal electron acceptor, reducing ATP production to its minimal level (no OXPHOS activity). *PDH* loss, either alone or in conjunction with other lost features, has a very limited impact on ATP production. Complete loss of the TCA cycle substantially compromises ATP production, but losses of many single steps (which would constitute *TCA* loss in our evolutionary analysis) have very little effect on ATP production; citrate synthase, CII, and malate dehydrogenase are exceptions where ATP production is more substantially reduced. The simulation under hypoxic conditions, and with proton-motive force rather than ATP generation as the metabolic objective have similar orderings of *CI, CIII, CIV*, and *TCA* features, with only differences in the relative importance of CV for the proton-motive force objective and CII for the hypoxic conditions (Supp. Fig. 5).

Once *CI* and any one of *CIII, CIV, TCA* are lost, ATP production approaches a minimum level. Hence, from an ATP production perspective, there is no advantage in retaining the other two members of the set. This compatible with the inference that loss of these features proceed in rapid succession in the CI-class pathway. Once either *CIII* or *CIV* is lost, loss of the other (or loss of many TCA steps) provides no further ATP production impact, again supporting the idea that this group of features can be lost in a highly coupled way.

Taking these observations together, there is a clear correlation between the modelled impact of feature loss on ATP production and the propensity of that feature to be lost early in our inferred evolutionary pathways. As such, for circumstances where ATP production remains an important component of fitness, the CI- and alt-class pathways may then reflect the smoothest routes of progress through intermediate states on a fitness landscape (see Discussion).

## Discussion

We have shown that the phylogenetically diverse, parallel instances of reductive mitochondrial evolution can be explained by a relatively limited set of canalised evolutionary pathways – and that in turn, these pathways can in part be explained by the relative metabolic impacts of the different feature losses involved. In particular, mitochondrial features seem to be lost roughly according to the consequent impact on metabolism.

A connection between reductive evolution of mtDNA gene content and metabolic demands from organismal environment and ecology has been proposed (García-Pascual et al., 2022; Giannakis, Richards, et al., 2024; Veeraragavan et al., 2024). Broadly, organelles in organisms with metabolism influenced by strong environmental oscillations (day/night cycles, tidal cycles) are predicted to retain more genes locally, while organelles in more stable and/or less demanding environments are freer to transfer genes to the nucleus. This idea borrows strongly from the theory of colocation for redox regulation (CoRR), suggesting that retaining organelle genes supports local, rapid, individual control of organelle machinery (Allen, 2015; Allen & Martin, 2016; Allen & Raven, 1996).

In this picture, reduced pressure to retain local control of mitochondrial function relaxes the pressure to retain mitochondrial genes in mtDNA, and favours their relocation to the safer nucleus. While many other factors are likely involved in shaping the reduction of mtDNA and mitochondria across species (Butenko et al., 2024; Johnston & Williams, 2016; Lynch et al., 2006; Smith, 2016), the loss of the features we consider here can in a sense be viewed as an extension of this picture, where the function of some mitochondrial machinery is under so little retention pressure that it is lost entirely. The properties of MROs have indeed been cited as evidence for CoRR over competing hypotheses for gene retention (de Paula et al., 2012).

Evolutionary accumulation modelling (EvAM) is at the core of our analysis here. EvAM approaches have often been applied only to cross-sectional observations, largely to study cancer progression in independent patients (Diaz-Uriarte & Johnston, 2024). EvAM approaches supporting phylogenetically-embedded data (Aga et al., 2024; Greenbury et al., 2020; Johnston & Williams, 2016; Luo et al., 2023; Moen & Johnston, 2023) have started to bridge the gap with more established tools from phylogenetic comparative methods (Boyko & Beaulieu, 2021; O’Meara, 2012; Pagel, 1994), with recent work explicitly phrasing EvAM as a well-known Mk model (Johnston & Diaz-Uriarte, 2024) and many EvAM approaches falling into such a continuous-time Markov chain with finite state space (CTMC-FSS) picture (O’Meara, 2012). We hope that this study illustrates that EvAM can provide insights into evolutionary systems involving the coupled evolution of multiple binary traits, complementing its existing use cases from C_4_ photosynthesis (Williams et al., 2013) to tool use in animals (Johnston & Røyrvik, 2020).

Our compilation and analysis of data from diverse eukaryotes is subject to several potential issues. Even the primary source data for the presence and absence of mitochondrial features is subject to some uncertainty (for example, when a feature is lost but its genes remain, or when a feature remains but is not detectable in particular experiments); the choices made in our binarisation of that source data may also be questioned. Species names and taxonomic positioning may also be updated over time – for example, since beginning this manuscript we have learned of a proposed reclassification for *Sawyeria marylandensis* into genus *Psalteriomonas* (Foučková et al., 2024). Some concerns about the robustness of our findings to data errors and changes may be resolved by the fact that introducing random errors in the source data did not qualitatively change the outcomes (Supp. Fig. 2), but we would always appreciate corrections for any obvious errors in our compilation. Further, in our effort to be general we have omitted specific details and features. The diversity of compensatory, non-canonical features gained by reduced mitochondria is not captured in our coarse-grained metabolic modelling; we do not consider transporters (like AAC), import machinery, the presence of other metabolic pathways, or other interesting MRO features in our dataset (Makiuchi & Nozaki, 2014). A clear target for future work is to expand this feature set, building on careful characterisation of these features across eukaryotes, which will increase the detail with which these convergence reduction pathways can be described.

## Acknowledgments

We are very grateful to Hassan Hashimi, Vladimír Hampl, Tomáš Pánek, Ingrid Sveráková, and other delegates of the 54th Jírovec’s Protozoological Days conference for feedback on the data and inferences in this study, and to Ellen Røyrvik, Paul Una, Amelia Earl, and Ai Monti for valuable discussions. This project has received funding from the European Research Council (ERC) under the European Union’s Horizon 2020 research and innovation programme (Grant agreement No. 805046 (EvoConBiO) to IGJ). This work was supported by the Trond Mohn Foundation [project HyperEvol under grant agreement No. TMS2021TMT09 to IGJ], through the Centre for Antimicrobial Resistance in Western Norway (CAMRIA) [TMS2020TMT11].

## Supplementary Information

**Supplementary Figure 1.**
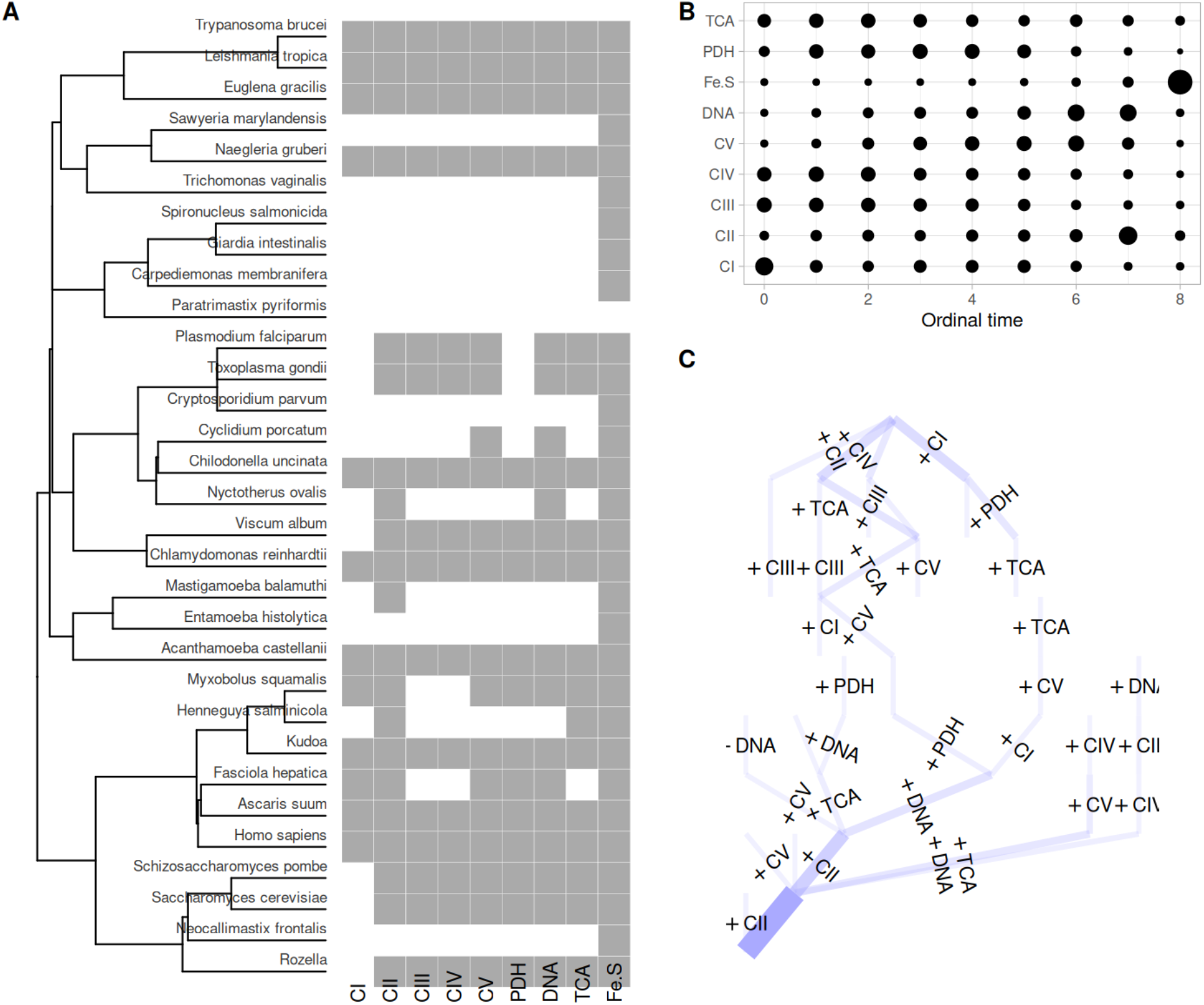
HyperTraPS-CT inference using continuous time. **(A)** Source data, with branch lengths now derived from divergence time estimates with TimeTree (Kumar et al., 2022). For scale, the top two species (*T. brucei* and *L. tropica*) have a divergence time of 266 Ma. **(B)** Bubble plot and **(C)** transition network from HyperTraPS-CT inference, now using a possible time window for each transition of width [0.5*τ*, 1.5*τ*] where *τ* is the estimated divergence time. Timescale distributions for these pathways are shown in Supp. Fig. 4.

**Supplementary Figure 2.**
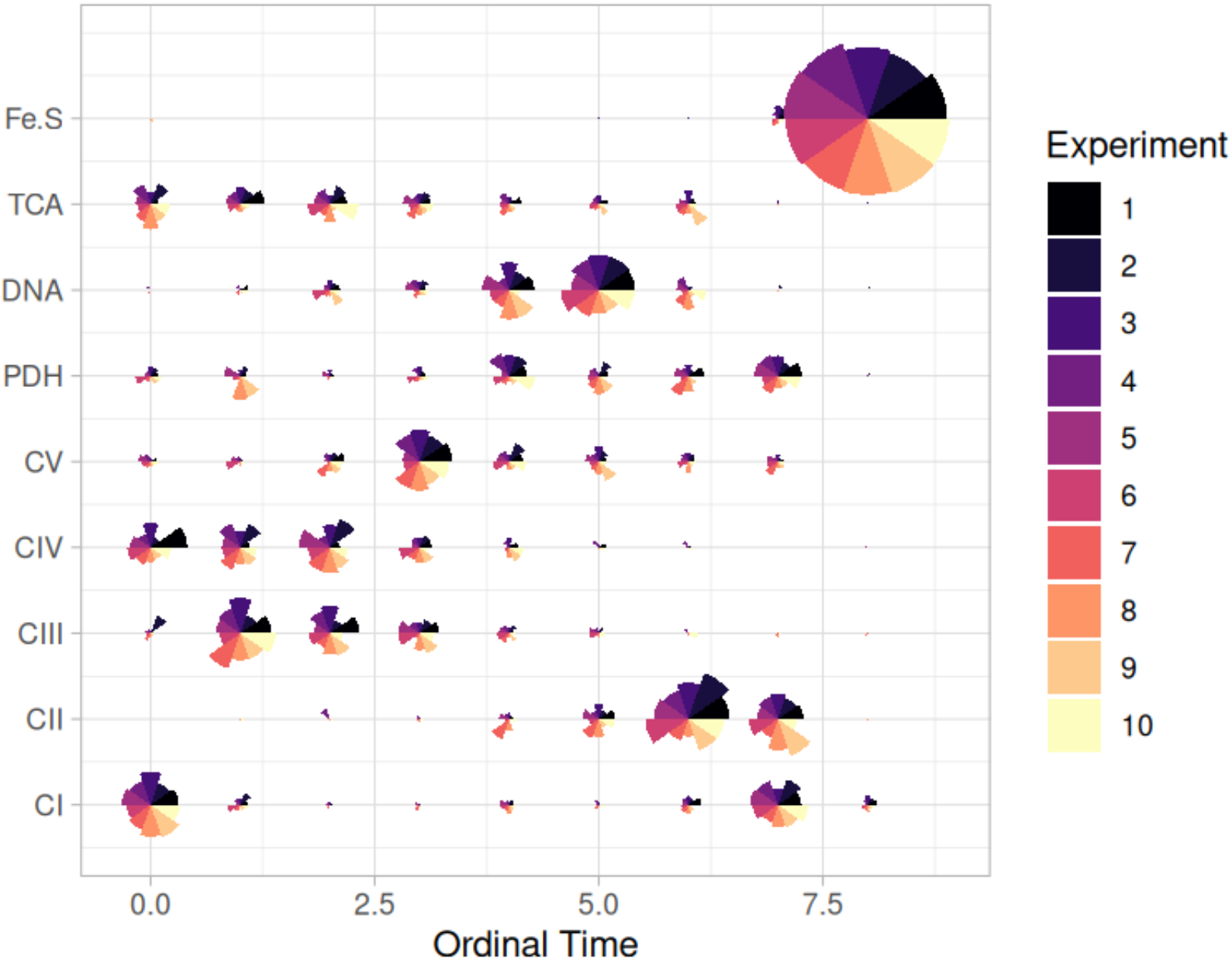
Robustness of inferred pathways with respect to uncertainty in source data. Bubble plot from HyperTraPS inference. Each segment represents the inference from a resampled dataset, where 10% of the data are artificially changed from 0 to 1 or vice versa. Overall the inferred dynamics are robust to these sets of perturbations, with segments showing the same trends across resamples.

**Supplementary Figure 3.**
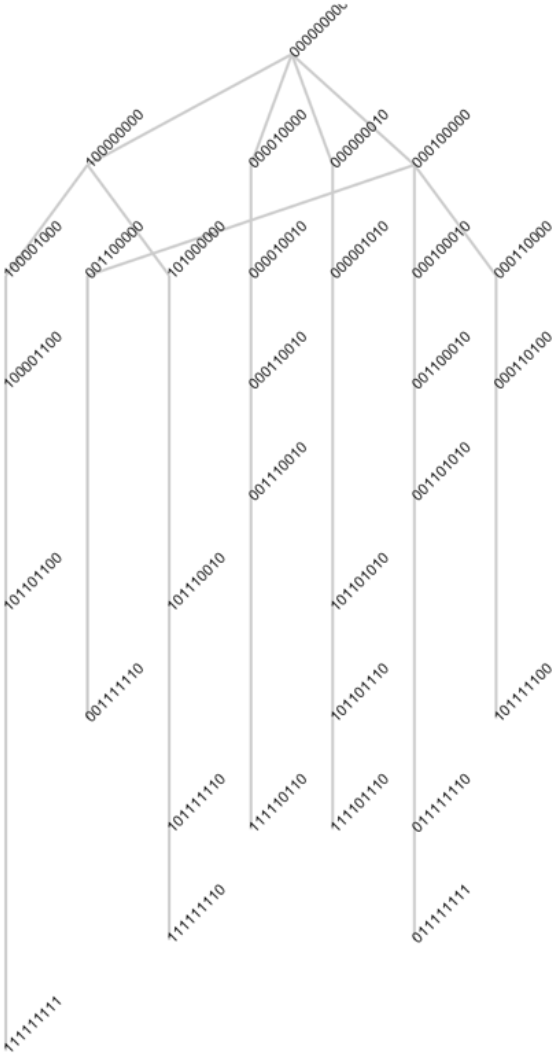
Hypercubic directed acyclic graph (HyperDAG) analysis of minimal pathways. A set of edges on the transition network that can account for all observed states in the source data (Fig. 1A) and display minimal branching – keeping as close as possible to a single, deterministic evolutionary pathway (Giannakis, Aga, et al., 2024). Each node is labelled by a binary string describing the features at which losses have (1) or have not (0) occurred: the ordering of features in these strings is *CI, CII, CIII, CIV, CV, PDH, TCA, DNA, TCA, Fe-S*. The top state, as in Fig. 1C, is the ancestral state with no losses. There is then a set of branches to *CI* (1000…), *CIII, CIV, TCA* first losses. The branches from *CIV* first loss (00010…) account for different patterns of the early alt-class steps. The branch from *CI* first loss accounts for either *CIII* or *PDH* being the next loss. Apart from these branches, all other states can be accounted for by canalised pathways, which descend the remainder of the evolutionary space.

**Supplementary Figure 4.**
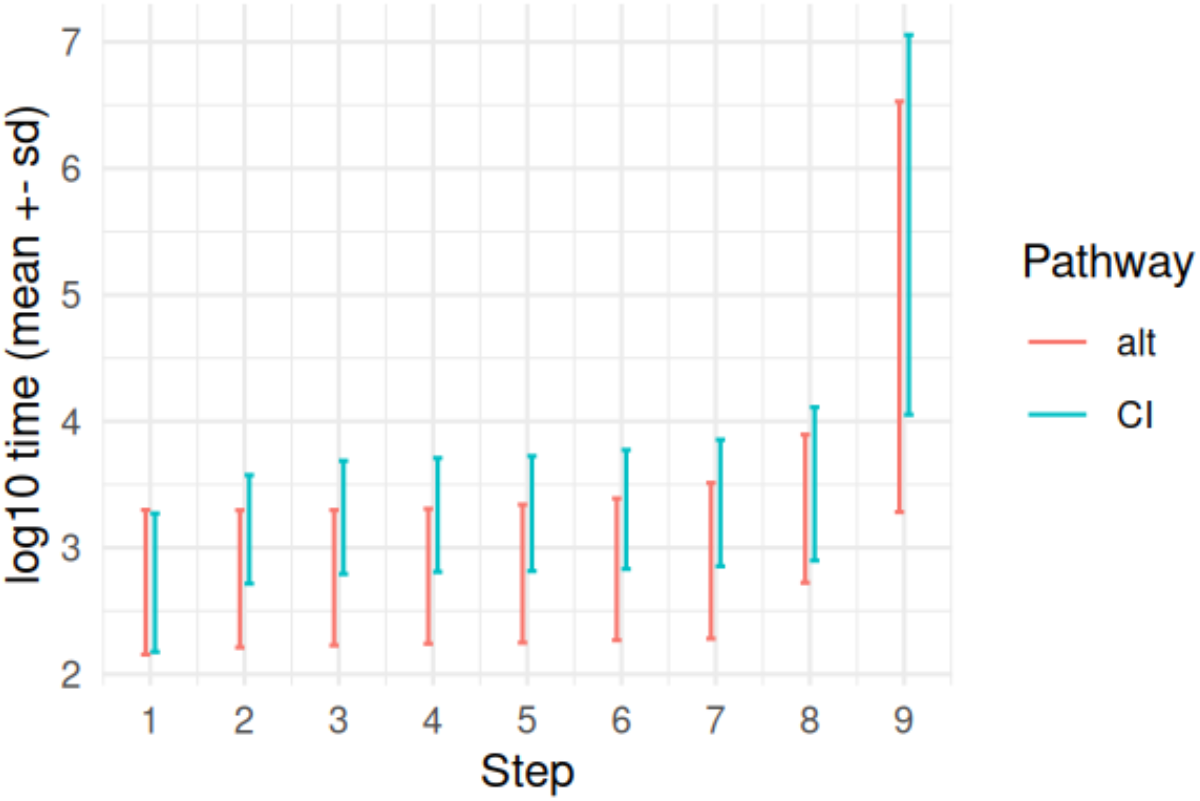
Inferred timescales of loss events in CI- and alt-class pathways. HyperTraPS-CT infers the distribution of timescales for each loss event in a model evolutionary process. The bars here give (mean ± standard deviation) for (log) timings of each step in a set of simulated CI-class and alt-class pathways on the fitted hypercubic model. For example, the first step for CI has a range of timings of 150-1900 Ma.

**Supplementary Figure 5.**
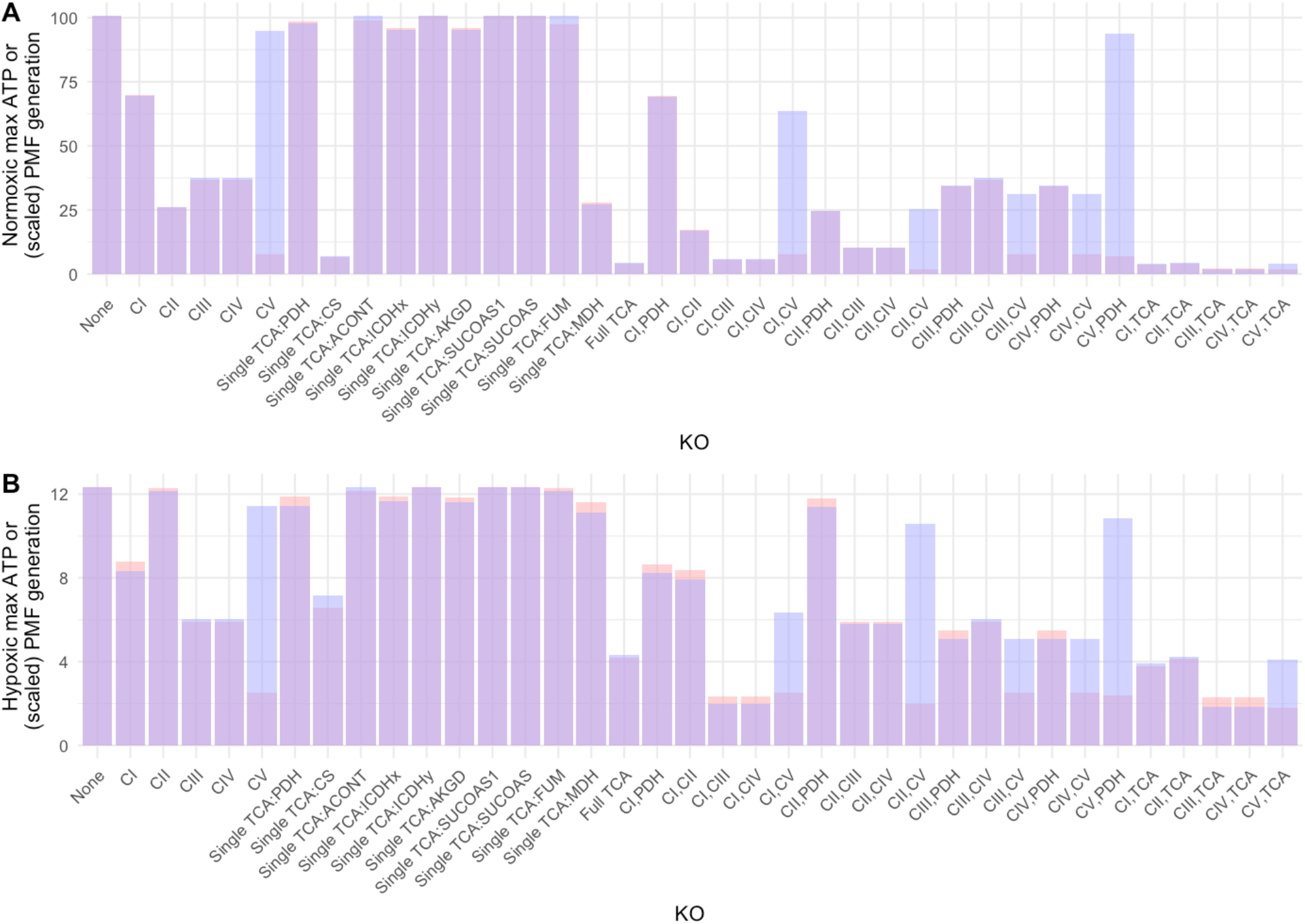
Flux balance analysis of mitochondrial feature loss under different conditions. (A) Normoxic and (B) hypoxic (10% normoxic) conditions simulated with simulated knockouts (KO) of different individual and paired mitochondrial features, as in Fig. 4. Red bars give flux values for ATP production under an objective of maximising ATP production, blue bars give proton motive force (PMF) under an objective of maximising PMF. PMF values are scaled by 1/3.7 to align with ATP values in the default model case with no knockouts.

Supplementary Text 1. **Additional references used in the meta-analysis of mitochondrial feature profiles**. Numbers correspond to labels in the annotated dataset.

